# Effects of expectation, attention, and NMDA receptor blockade on feedforward and feedback processing

**DOI:** 10.64898/2026.02.25.707924

**Authors:** Samuel Noorman, Johannes J. Fahrenfort, Micha Heilbron, Claire Sergent, Jasper B. Zantvoord, Simon van Gaal, Timo Stein

## Abstract

Perception is increasingly viewed as an inferential process wherein sensory inputs are integrated with prior expectations. We employed time-resolved decoding on electroencephalography (EEG) data (n = 30 male participants) to investigate how expectations modulate sensory processing across varying levels of stimulus complexity, and tested the effect of attention and NMDA receptor blockade. We designed a visual stimulus containing features of different complexity whose processing relies on distinct neural mechanisms: local contrast, collinearity, and the Kanizsa illusion, involving primarily feedforward, lateral, and feedback processes, respectively. EEG decoding revealed that expectations modulated lateral and feedback processing (better decoding for unexpected stimuli) but not feedforward processing. These expectation effects were confined to attended (task-relevant) features and were not observed for task-irrelevant features. The NMDA receptor antagonist memantine selectively enhanced decoding of the Kanizsa illusion, implicating NMDA-mediated feedback mechanisms in perceptual inference, but it did not modulate the effects of expectation or attention. These findings highlight the differential impact of expectations across different stages of sensory processing and reveal a distinct role of NMDA receptor-mediated feedback mechanisms.

**Significance statement:** Perception integrates sensory inputs with prior expectations. Using EEG decoding, we examined how expectations shape sensory processing at different levels of complexity and tested the effects of attention and NMDA receptor blockade. Our visual stimuli were designed to capture EEG markers of feedforward, lateral, and feedback mechanisms. Expectations influenced lateral and feedback processing (better decoding for unexpected stimuli) but not feedforward processing, and these effects were selective to task-relevant stimulus features. Memantine, an NMDA receptor antagonist, selectively improved decoding of the Kanizsa illusion, implicating NMDA-mediated feedback in perceptual inference, but it did not shape expectation or attention effects.

## Introduction

Perception is widely hypothesized to be an inferential process in which bottom-up sensory input is combined with top-down expectations based on a generative model. Expectations can reflect long-term priors about contextual regularities, as well as short-term prior learned from stimulus probabilities (Bar, 2009; de Lange et al., 2018; Friston, 2005; Summerfield & de Lange, 2014). Prior expectations are passed down the processing hierarchy via feedback connections to earlier cortical areas, where they modulate responses to incoming sensory information. While this general framework is supported by extensive work (for review see de Lange et al., 2018; Walsh et al., 2020), we here address four central unresolved questions.

First, do expectations influence all processing stages equally, including early sensory stages (Kok, Jehee, et al., 2012; Murray et al., 2002; Richter & de Lange, 2019; Richter et al., 2017; Stein et al., 2022), or primarily later stages involved in processing more complex stimulus features (Alilović et al., 2019; Rungratsameetaweemana et al., 2018; Rungratsameetaweemana & Serences, 2019)? Second, given the established role of NMDA receptors in recurrent processing (Meuwese et al., 2013; Noorman et al., 2025b; ; Sachidhanandam et al., 2013; Self et al., 2012; Thiele, 2012; Thiele & Bellgrove, 2018; Van Kerkoerle et al., 2014; van Loon et al., 2016; Wang, 2008), do expectations involve NMDA-dependent recurrent processing? Third, do expectations enhance expected sensory inputs akin to attention enhancing task-relevant inputs, or do expectations suppress relatively uninformative sensory inputs (Press et al., 2020)? Fourth, do expectation effects occur automatically (den Ouden et al., 2012; Kok, Rahnev, et al., 2012; Richter et al., 2017; Stefanics et al., 2019), or do they require attention (Richter & de Lange, 2019), and does this depend on stimulus complexity?

To answer these questions, we used time-resolved EEG decoding in a full-factorial experimental design manipulating expectation and attention to measure processing across hierarchical stages (**Fig. 1A**). The stimulus dimensions differed systematically in complexity (a) local contrast, manipulated by a 180° rotation of the entire stimulus, which can be encoded via a single feedforward sweep (Fahrenfort et al., 2007, 2017; Kandel et al., 2000; Lamme & Roelfsema, 2000); (b) collinearity, defined by the alignment of three “two-legged” white circles to form a non-illusory (amodally completed; Pessoa et al., 1998) triangle, which depends primarily on lateral connections (Bosking et al., 1997; Gilbert & Wiesel, 1979; Li, 1998; Liang et al., 2017; Schmidt et al., 1997; Stettler et al., 2002); and (c) illusory Kanizsa triangle, induced by three aligned Pac-Man inducers, which additionally requires top-down feedback to generate a perceptually filled-in surface (Halgren et al., 2003; Kok et al., 2016; Kok & de Lange, 2014; Lee & Nguyen, 2001; Pak et al., 2020; Wokke et al., 2013). Expectations were induced by manipulating the probability of the task-relevant feature (75% vs. 25% of trials) within a block, akin to an oddball design. Attention was manipulated by designating a single feature as task-relevant per block.

**Figure 1.**
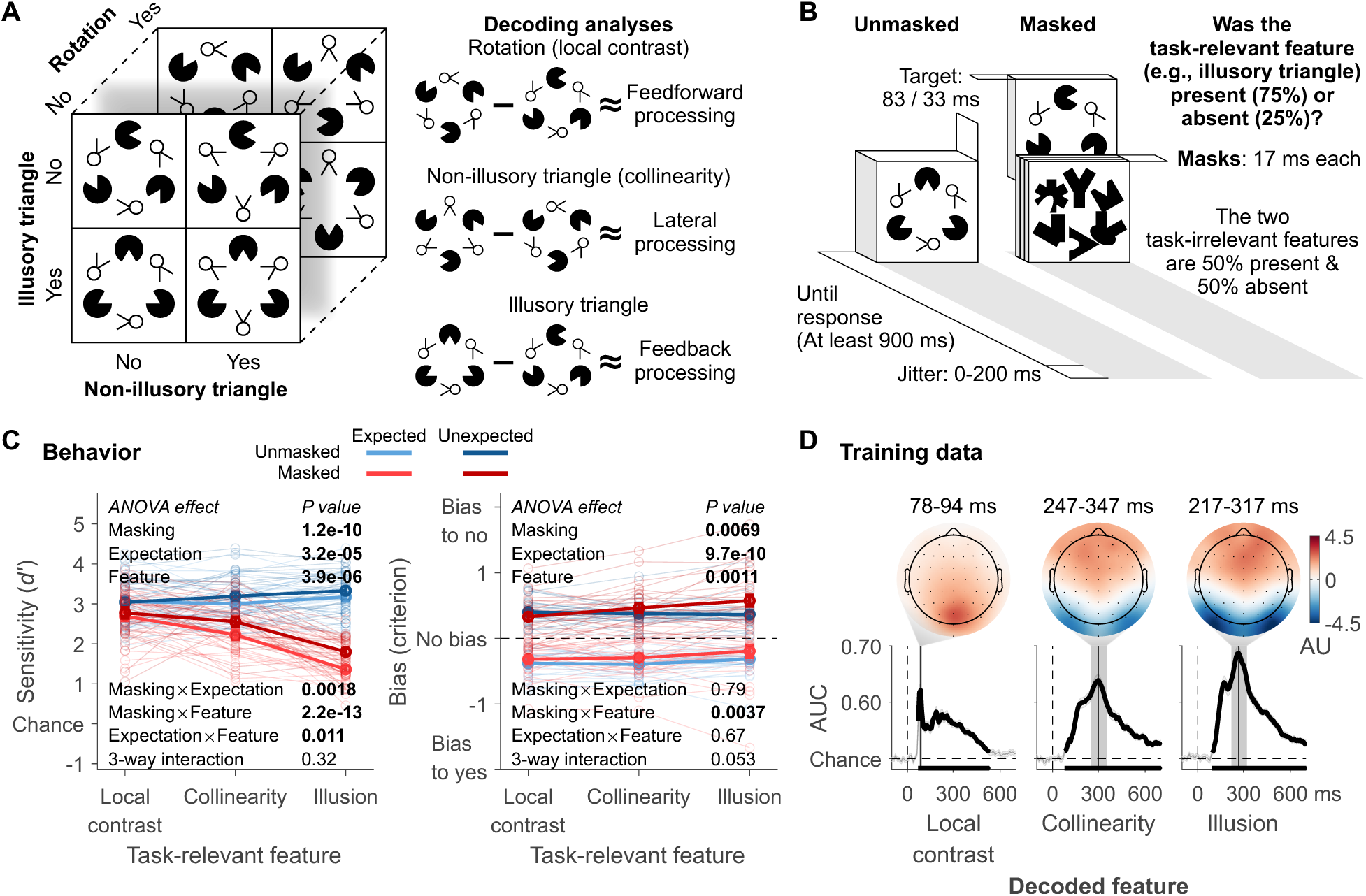
Experimental design, behavior and training data of the decoder. (A) Target stimulus set and schematic of the markers for the different types of processing: local contrast decoding (supported by feedforward connections), collinearity (forming an amodally completed, non-illusory triangle) decoding (supported by lateral connections), and illusory triangle (a modally completed, perceptually filled-in surface) decoding (supported by lateral and feedback connections) (B) Trial design. (C) Perceptual performance and bias related to participants’ ability to detect the task-relevant (attended) stimulus dimension. Error bars are mean ± standard error of the mean. Individual data points are plotted using low contrast. For completeness, behavioral performance for drug and placebo conditions is shown separately in Supplementary **Fig. S1**; no significant effect of drug or drug-related interactions were observed. (D) Upon testing the classifiers, each decoded visual feature exhibited a distinct peak in classifier performance (area under the receiver operating characteristic curve [AUC]) over time: local contrast at 86 ms after stimulus onset, collinearity at 297 ms, and the illusion at 266 ms). For all features, covariance/class separability maps reflecting underlying neural sources are shown. Thick black lines differ from chance: P<0.05, cluster-based permutation test.

To test whether expectation effects rely on NMDA-receptor mediated recurrent processing, we administered the NMDA antagonist memantine. Previous results suggested that memantine specifically modulates illusion decoding (Noorman et al., 2025a). Illusion processing can be regarded as involving perceptual inference based on long-term priors (Kok & de Lange, 2014; but see Moors, 2015), where the alignment of the inducers is expected (to form a “good Gestalt”; **Fig. 1A**). The key question was how neural effects of short-term expectations based on the manipulation of stimulus probability would be affected by memantine, and how such modulation would depend on feature complexity and attention. A backward-masking condition was included to ensure that the three features reflected distinct neural processes. Masking was expected to interfere with recurrency-dependent collinearity and illusion decoding, while leaving feedforward-mediated local contrast decoding largely intact (Dehaene et al., 2006; Fahrenfort et al., 2007, 2017; Lamme, 2010; Lamme et al., 2002; Noorman et al., 2025a, 2025b).

## Methods

### Participants

Thirty right-handed men with normal, or corrected-to-normal, vision were recruited after they had passed extensive physical and mental screening. Two of them dropped out after the first experimental session because of personal reasons unrelated to memantine. EEG recording failed during an experimental session of one participant. The data from the remaining 27 participants (22±2 years old) were analyzed. The study was approved by the Medical Ethical Committee of the Amsterdam University Medical Centre (AUMC) (NL64341.018.18) and the local ethics committee of the University of Amsterdam (2022-BC-14355). Participants gave informed consent and received monetary compensation.

### Stimuli

The target stimulus set had a 2 Kanizsa illusion levels (present/absent) × 2 collinearity levels (present/absent) × 2 rotation (local contrast) levels (present/absent) design, resulting in eight stimuli (**Fig. 1A**). Importantly, the three stimulus dimensions were fully orthogonalized: the presence or absence of each feature (local contrast, collinearity, illusion) varied independently and occurred equally often across conditions. This orthogonalization prevents systematic dependencies between features, ensuring that any task-irrelevant feature did not consistently co-occur with, or depend on, any other feature. As a result, potential higher-order feature interactions (e.g., whether detecting the non-illusory triangle is more difficult when an illusory triangle is also present) could not bias the decoding analyses. As described in the Introduction, the three stimulus features were designed to preferentially (but not uniquely) engage different stages of visual processing: local contrast primarily involves feedforward processing, collinearity engages primarily lateral recurrent interactions, and the Kanizsa illusion additionally recruits top-down feedback recurrence.

Three aligned Pac-Men induced the Kanizsa illusion, generating a perceptually filled-in (modally completed) triangle. Collinearity was present when the stimuli’s three other elements (the “two-legged white circles”) were aligned to form a non-illusory (amodally completed) triangle. Local contrast differences were created by rotating the entire stimulus 180 degrees, reversing the polarity of black and white regions. Controls for the illusion and collinearity were created by rotating their respective elements (Pac-Man and two-legged white circle stimuli) by 90°.

The targets spanned 7.5 degrees by 8.3 degrees of visual angle. The distance between the three Pac-Man stimuli as well as between the three aligned two-legged white circles was 2.8 degrees of visual angle. Although neuronal responses to collinearity in primary visual cortex are most robust when this distance is smaller (Kapadia et al., 1995, 2000), longer-range horizontal connections between neurons with similar orientation selectivity can span distances corresponding to visual angles considerably greater than 2.8 degrees (Bosking et al., 1997; Stettler et al., 2002).

Masks consisted of six differently shaped elements, all capable of covering the targets’ elements. Six masks were created by rotating the original mask five times by 60 degrees. They spanned 8.5 degrees by 9.1 degrees visual angle. The design of the fixation cross, which was present throughout the trial, was adapted from Thaler et al. (2013).

### Task designs

#### Localizer task

Decoding analyses were based on an independent training data set. The three different visual features of the target stimulus (Kanizsa illusion, collinearity, rotation) were task-relevant (attended) in separate blocks. On each trial, a target stimulus was presented for 83 ms.

Participants fixated on the fixation cross and indicated whether the current task-relevant feature (i.e., the Kanizsa illusion, the non-illusory triangle, or the downward pointing stimulus) was present or absent. The (first) response window started at target onset and ended after 900 ms. If no response was made, “Respond faster!” was presented until response. Trials ended with 0-200 ms jitter and at least 300 ms between response and jitter. The order of the task-relevant features was counterbalanced over participants and, because each participant performed the task multiple times, kept the same within participants. Response button mapping was counterbalanced within the task. Each localizer task consisted of 1728 trials, evenly distributed across 12 blocks. Participants completed the task three times, in each separate session (intake, and two main sessions), for a total of 5184 trials. In the intake, no drugs were administered. In the other two sessions the localizer task was performed well before memantine reached peak plasma levels (directly after pill intake when memantine was not yet or barely active).

#### Main task

The main task (**Fig. 1B**) was similar to the localizer task with some crucial additions.

To manipulate stimulus visibility, half the trials were masked. On these trials, the target stimulus was presented for 33 ms and immediately followed by three masks, presented for 17 ms each. The three masks were selected at random with the constraint that all differed from each other.

To manipulate attention, similar to the localizer, each different visual feature of the target stimulus (Kanizsa illusion, collinearity, rotation) was task-relevant (attended) in some blocks and task-irrelevant in other blocks. Participants performed 4 blocks of each task-relevance condition.

To manipulate expectation, the task-relevant feature was made high-probability in one block (present on 75% of trials) and low-probability in another block (present on 25% of trials), following an oddball-like base-rate manipulation that is widely used to study perceptual expectations (Feuerriegel et al., 2021). The order of these two block types was counterbalanced across participants, and participants were informed about the upcoming block type. In this design, expectation for the task-relevant feature is defined as an expectation about its presence versus absence.

Expectation effects for task-relevant features were quantified by decoding feature presence versus absence within each block. For example, in a block where the Kanizsa illusion was high-probability, the illusion was present on 75% of trials and absent on 25%; the “expected” condition contrasted illusion-present versus illusion-absent trials from that same block. Conversely, in a block where the illusion was low-probability, it was present on 25% of trials and absent on 75%, and the “unexpected” condition was defined using the same within-block presence-versus-absence contrast. Thus, decoding of task-relevant features was always performed within blocks, and expectation effects reflect differences between high- and low-probability outcomes under a constant block context.

Task-irrelevant features were always equiprobable (50% present, 50% absent) and rarely repeated on consecutive trials. As a result, no direct expectation about the presence or absence of task-irrelevant features was induced. For task-irrelevant features, expectation was therefore defined with respect to the probability of the task-relevant feature with which they co-occurred, rather than the probability of the task-irrelevant feature itself.

Decoding of task-irrelevant features followed the same general logic as for task-relevant features. Presence versus absence of the task-irrelevant feature was decoded depending on whether it was presented as part of a high-probability (75%) or low-probability (25%) task-relevant outcome. For example, when decoding collinearity in blocks where the Kanizsa illusion was task-relevant, collinearity presence versus absence was first decoded separately within each block type, using illusion-present trials in blocks with 75% illusion presence and illusion-absent trials in blocks with 25% illusion presence (75% illusion absent). The resulting within-block decoding estimates were then combined to define the *expected* task-irrelevant condition. Analogously, for the *unexpected* task-irrelevant condition, collinearity presence versus absence was decoded within blocks using illusion-present trials in blocks with 25% illusion presence and illusion-absent trials in blocks with 75% illusion presence (25% illusion absent), and these within-block decoding estimates were combined. Thus, task-irrelevant features co-occurring with a high-probability task-relevant outcome were treated as “expected,” regardless of whether that outcome involved presence or absence of the task-relevant feature. Thus, expectation effects on decoding of a task-irrelevant feature reflected whether presence or absence of the task-relevant feature was high- or low-probability in that block.

To manipulate NMDA-receptor functioning, Memantine (20 mg) and placebo were administered in different experimental sessions/days, separated by at least a week (randomized, double-blind crossover design; more details in the Procedure section below).

In summary, the task design had 2 masking levels (present/absent) × 3 task-relevance conditions (rotation, non-illusory triangle, illusory triangle) × 2 probability rates for the task-relevant feature (75%/25%) × 2 drug conditions (memantine/placebo). In total there were 7680 trials in the main task.

### Procedure

The experiment consisted of three separate sessions conducted on different days: a four-hour screening session and two seven-hour experimental sessions. At the beginning of the screening session, the volunteers were screened for contraindications to using memantine. If there were no contraindications, participants proceeded to performing the task for the localizer (independent training data set) and practicing the main task.

The practice session included the staircase procedure to determine the mask strength for subsequent sessions. We aimed for a masked performance of ∼75% correct by staircasing mask contrast using the weighted up-down method (Kaernbach, 1991). Staircasing was done when the Kanizsa illusion was task-relevant, because masking affected this feature more than the other two. Contrast levels ranged from 0 (black) to 255 (white). Mask contrast started at level 230. Each correct response made the task more difficult: masks got darker by downward step size *S*_down_. Each incorrect response made the task easier: masks got lighter by upward step size *S*_up_. Step sizes were determined by *S*_up_ × *p* = *S*_down_ × (1 - *p*), where *p* is the desired accuracy 0.75. The smallest step size was always 13 contrast levels. A reversal is making a mistake after a correct response, or vice versa. The staircase ended after 25 reversals. The mask contrast that was used during the experimental sessions was the average contrast level of the last 20 reversals.

Memantine (20 mg) and the placebo were administered in different experimental sessions, separated by at least a week. These sessions were conducted using a randomized, double-blind crossover design. The order of drug administration (placebo or memantine first) was counterbalanced across participants. Participants were required to refrain from recreational drugs for four weeks before the experiment and to refrain from alcohol for two days before each experimental session.

Experimental sessions started at 9:00 with the pill intake, which was immediately followed by another performance of the task for the independent training data set to increase the power of the decoding analyses. This was followed by briefly practicing the main task (576 trials).

The experimental sessions of the main task were performed when memantine reached peak plasma level, four hours after the pharmaceutical was administered. Before pill intake (∼9:00) and before (∼13:00) and after (∼15:30) the main task, participants filled in the visual analogue scale (VAS) to measure the effects of memantine on subjective state (Bond & Lader, 1974; Danion et al., 1989). The mean scores of subsets of the VAS were calculated and taken as measures of alertness, contentedness, and calmness, following Bond and Lader (1974), and as a measure of sedation, following Danion et al. (1989). VAS scores of the two measurements after pill intake were averaged and corrected by calculating percentage change from the baseline measurement. One of the baseline VAS forms of one participant was not filled in, so this participant was excluded from the VAS analysis. VAS measurements revealed no side effects from memantine (all four measure’s BF_01_>3.59).

Tasks were programmed in Presentation software (Neurobehavioral Systems). For 18 participants, tasks were displayed on a 23 inch, 1920 × 1080 pixels monitor and for the other participants on a 27 inch, 2560 × 1440 pixels monitor, but stimulus size was the same in the two different labs. The monitor was viewed from a distance of 80 cm distance with the use of a chin rest.

### Behavioral analysis

To investigate behavioral performance in the main task, we used perceptual sensitivity (d’) and bias (criterion) derived from signal detection theory. Repeated measures ANOVAs and paired *t*-tests were used to test for differences between conditions.

### EEG recording and preprocessing

EEG was recorded at 1024 Hz using a 64 channel ActiveTwo system (BioSemi). Four electrooculographic (EOG) electrodes measured horizontal and vertical eye movements. The data were analyzed with MATLAB (MathWorks). For most of the preprocessing steps, EEGLAB was used (Delorme & Makeig, 2004). The data were re-referenced to the earlobes. Poor channels were interpolated. High-pass filtering can cause artifacts in decoding analyses, so for slow-drift removal we used trial-masked robust detrending (van Driel et al., 2021). Each target was epoched from −250 to 1000 ms relative to target onset. To improve the independent component analysis (ICA), baseline correction was applied using the whole epoch as baseline (Groppe et al., 2009). ICA was used to remove blinks. Blink components were removed manually. Baseline correction was applied, now using a −250 to 0 ms window relative to target onset. Trials with values outside of a −300 to 300 microvolts range were removed. We used an adapted version of FieldTrip’s ft_artifact_zvalue function to detect and remove trials with muscle artifacts (Oostenveld et al., 2011). Finally, the data were downsampled to 128 Hz.

### Multivariate pattern analysis

We used the Amsterdam Decoding and Modeling (ADAM) toolbox to perform multivariate pattern analyses (MVPAs) on the EEG data (Fahrenfort et al., 2018). Using a tenfold cross-validation scheme, we first analyzed the data from the independent training set. We decoded each visual feature (e.g., the Kanizsa illusion) by training linear discriminant classifiers to discriminate between the feature’s presence and absence based on the preprocessed EEG data. Only trials during which the decoded feature was task-relevant were included. Individual participants’ data were split into ten equal-sized folds after randomizing trial order. A classifier was then trained on nine folds and tested on the tenth one, ensuring independence of the training and testing sets. This procedure was repeated until each fold served as the test set once. Classifier performance, here area under the receiver operating characteristic (AUC) (Hand & Till, 2001), was averaged across all ten iterations. Classifier performance was calculated for every time sample and the classifiers were tested on the same time sample they were trained on. To visualize how representations evolved and generalized over time, we also computed full temporal generalization matrices (King & Dehaene, 2014), training classifiers on each time point and testing them on all other time points. These matrices, shown in the Supplementary **Fig. S2**, confirm that the temporal dynamics of decoding were stable across conditions and that our main diagonal-based approach captured the peak information content within each feature’s relevant time window.

Each decoded visual feature showed a clear peak in classifier performance over time (**Fig. 1D**). The ANOVAs are based on the averages of the time windows encompassing these peaks, similar to our previous studies (Fahrenfort et al., 2017; Noorman et al., 2025a, 2025b): from 78 to 94 ms for local contrast, 247-347 ms for collinearity, and 217-317 ms for the illusion. These peak-based windows were chosen as feature-specific, data-driven markers of the dominant representational phase for each feature. All three features showed earlier onsets of decodability, but collinearity and illusion exhibited additional later peaks, which we interpret as reflecting relatively stronger contributions from recurrent processing. Using fixed-latency or onset-based windows instead would confound differences in representational strength with differences in timing across features, whereas peak-based windows isolate the epochs where each feature is most strongly represented. To obtain topographic maps showing the neural sources of the classifier performance, we multiplied the classifier weights with the data covariance matrix, yielding covariance/class separability maps (Haufe et al., 2014). The covariance/class separability maps of our three time windows show that the classifiers depended on occipitoparietal electrodes during peak decoding accuracy (**Fig. 1D**). We restricted our decoding analyses to using only these electrodes (TP7, CP5, CP3, CP1, CPz, CP2, CP4, CP6, TP8, P9, P7, P5, P3, P1, Pz, P2, P4, P6, P8, P10, PO7, PO3, POz, PO4, PO8, O1, Oz, O2, and Iz) (see also Fahrenfort et al., 2017; Noorman et al., 2025b) to ensure that any effects we observed were not due to poor signal-to-noise ratio.

For the main analyses, classifiers were trained on the independent training set’s data and tested on each condition of the main task’s data. Classifier performance was calculated for every time sample and the classifiers were tested on the same time sample they were trained on. For each visual feature, we then averaged decoding within the feature-specific time window that had been identified in the training set (i.e., the time window in which that feature was decodable in the localizer). This ensured that the same, independently defined temporal window was used when quantifying decoding in the main task. To establish the markers for the different forms of neural processing, each visual feature was decoded in this manner. As for the behavioral analyses, repeated measures ANOVAs and paired *t*-tests—here applied to the mean decoding within these predefined time windows—were used to test for differences between conditions.

For the auxiliary analyses of decoding time courses, we analyzed time samples from −100 to 700 ms relative to target onset using a two-sided *t*-test to evaluate whether classifier performance differed from chance. We used cluster-based permutation testing (10,000 iterations at a threshold of 0.05) to correct for multiple comparisons (Maris & Oostenveld, 2007).

To assess whether our training procedure could bias the difference between task-relevant and task-irrelevant decoding, we performed a control analysis in which classifiers were trained (1) only on task-relevant localizer data (as in the main analysis), (2) only on task-irrelevant localizer data, and (3) on both combined. The resulting decoding patterns were qualitatively similar across all three training schemes, confirming that the observed task-relevance effects were not driven by classifier training biases (**Fig. S3**).

Control analyses tested whether expectation effects reflect adaptation related to feature repetition (Feuerriegel et al., 2021; Feuerriegel, 2024). We split trials into repeated versus non-repeated (same feature as previous trial vs. not) and assessed the distinct contribution of repetition and expectation on AUC within each decoding window (**Fig. S4**).

Note that the decoding analyses were not designed to predict perceptual reports on individual trials. Because each condition contained relatively few incorrect trials, single-trial decoding–behavior correlations were statistically underpowered and not theoretically expected in this paradigm (see also Noorman et al., 2025a).

### Data availability statement

Note that largely non-overlapping results from this dataset have been published elsewhere (Noorman et al., 2025b). In that paper we showed that the administration of memantine selectively enhanced decoding of the Kanizsa illusion but not decoding of local contrast and collinearity. In the present work we relate this finding to manipulations of task-relevance and expectation.

All code and data related to the present paper are available at UVA/AUAS figshare (we will add a link when the paper is accepted).

## Results

Twenty-seven participants performed a simple feature detection task in which each of three independently manipulated visual features (rotation [i.e., local contrast], non-illusory triangle [i.e., collinearity], or Kanizsa illusion) was task-relevant in a different block (attention manipulation, **Fig. 1B**). On each trial, participants indicated whether the task-relevant feature (e.g., the Kanizsa illusion) was present or absent. A target stimulus was presented for 83 ms when it was unmasked and for 33 ms when it was masked (see Methods for details). Expectations were induced by manipulating the probability of the task-relevant feature in different blocks of trials: the task-relevant feature was either present in 75% of the trials (e.g., illusion present) or in 25% of the trials. This constitutes an oddball design with a base-rate manipulation that has widely been used to study perceptual expectations (Feurriegel et al., 2021); critically, task-irrelevant features were equiprobable and seldom repeated, permitting tests of expectation effects beyond repetition. This was a pharmacological study with a randomized, double-blind, crossover design in which we administered the NMDA antagonist memantine in one session and a placebo in another session (on a different day).

### Behavioral results

Behavioral performance in the feature detection task was analyzed as sensitivity (ability to distinguish between feature presence and absence) and response criterion (tendency to indicate feature presence), following signal detection theory. The complete results from repeated-measures ANOVAs with the factors stimulus feature (local contrast, collinearity, illusion), expectation (expected, unexpected), masking (unmasked, masked) and drug (memantine, placebo) are shown in **Fig. 1C**. Here we focus on the most important findings.

Sensitivity was impaired by masking, as would be expected, and this effect was strongest for the illusion and weakest for local contrast. This suggests that the effect of masking increased with the requirement of recurrent processing for feature detection. Note, however, that the elements of the masks (see **Fig. 1B**) were more similar to the inducers of the Kanizsa illusory triangle (the “Pac-Man” stimuli) than to the inducers of the non-illusory triangle (the “two-legged white circle” designed to test for collinearity processing), which could explain the stronger effect of masking on illusion detection. Sensitivity was higher in blocks where the presence of a target feature was 25% likely (i.e., unexpected) than in blocks where it was 75% likely (i.e., expected), and this effect of expectation was specific to collinearity and the illusion. Turning to the effect of expectation on response criterion, subjects were more likely to report feature presence when the feature was expected than when it was unexpected. In contrast to sensitivity, this expectation effect was present for all three stimulus features, likely representing a general response strategy reflecting the differences in target probability.

For completeness, behavioral data are shown separately for drug and placebo conditions in Supplementary **Fig. S1**; consistent with the statistical analyses, no significant drug effects or interactions were present.

EEG decoding results described next showed the same qualitative pattern as the behavioral effects across conditions, e.g., stronger masking and expectation effects for collinearity and illusion than for contrast.

### Markers for feedforward, lateral, and feedback processing

To delineate EEG markers for the distinct neural processes of interest – feedforward, lateral, and feedback processing – we decoded the processing of various visual features – local contrast, collinearity, and the Kanizsa illusion – respectively (see **Fig. 1A**). Classifiers were trained using an independent data set acquired using a version of the experimental task without the conditions of masking, stimulus probability, and drugs. The time windows for feedforward (local contrast), lateral (collinearity), and feedback (illusion) processing were determined by decoding each visual feature from the independent training data set using a tenfold cross-validation scheme (see Methods). Each decoded visual feature exhibited a distinct peak in classifier performance (area under the receiver operating characteristic curve [AUC]) over time: local contrast at 86 ms after stimulus onset, collinearity at 297 ms, and the illusion at 266 ms.

Averages around these time points were used for follow-up ANOVAs, as in previous studies (see **Fig. 1D** for the exact time windows used) (Noorman et al, 2025a, 2025b). For the main analyses, classifiers trained on the independent training set were tested on each condition of the main task. The decoding time windows were defined feature-specifically, based on the temporal epochs in which each feature could be reliably decoded in the independent training data. We therefore use the terms ‘feedforward’, ‘lateral’, and ‘feedback’ in a relative sense to describe these empirically defined, feature-specific decoding time windows. Early “feedforward” windows likely already contain some recurrent activity, and “lateral” windows likely encompass both lateral and feedback processes and so forth, so the mapping should not be interpreted as strictly one-to-one between feature and processing stage.

### Masking leaves EEG markers of feedforward processing largely intact

We first focused on the effect of masking to confirm that decoding the different stimulus features indeed isolated different neural mechanisms. Masking is widely thought to impact later, recurrent stages of processing more strongly than earlier, predominantly feedforward stages.

Indeed, we found that masking impaired illusory and non-illusory (collinearity) triangle decoding much more than local contrast decoding (feature x masking interaction: *F*_2,52_=45.98, *P*<10^-9^, η^2^_p_=0.64). **Fig. 2A** shows the decoding time courses for each of the three different features.

**Figure 2.**
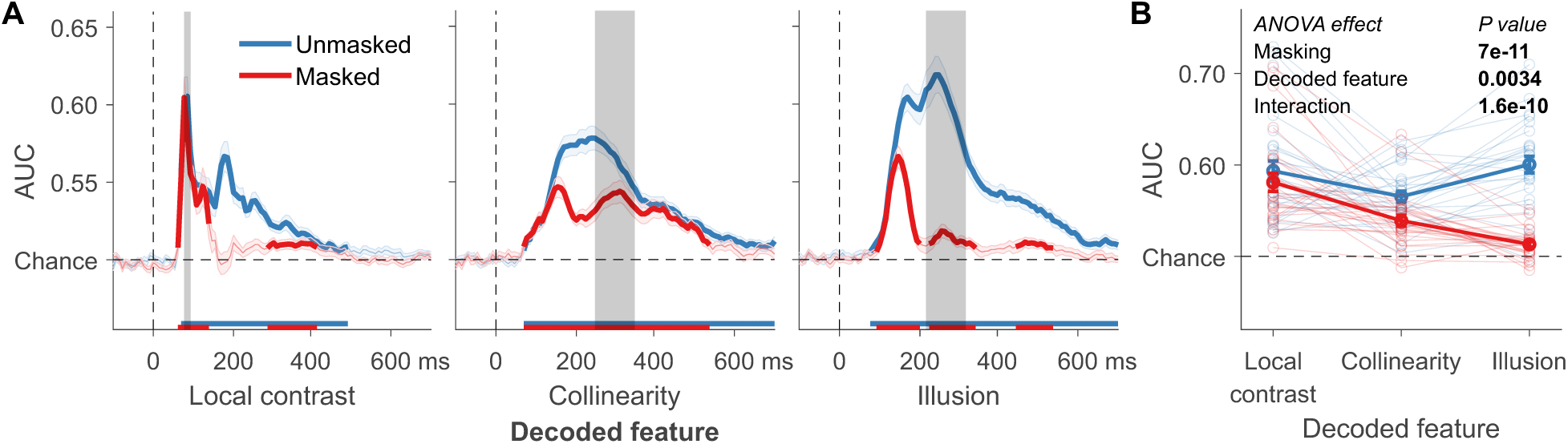
(A) The effect of masking on local contrast, non-illusory triangle, and illusory triangle decoding. Mean decoding performance, area under the receiver operating characteristic curve (AUC), over time ± standard error of the mean (SEM). Thick lines represent time-points that differ from chance: P<0.05, cluster-based permutation test. (B) Every time window’s mean AUC for every condition. Error bars are mean ± SEM. Individual data points are plotted using low contrast.

Although masking did have a small but significant effect on local contrast decoding (*t*_26_=4.60, *P*<0.001), this effect was much smaller than for the non-illusory and illusory triangle (effect sizes: Cohen’s d of 0.88 vs. 0.96 and 1.86, respectively, see **Fig. 2B**), consistent with the idea that masking preferentially disrupts later, predominantly recurrent processing stages relative to earlier, more feedforward-dominated processing (Dehaene et al., 2006; Fahrenfort et al., 2007, 2017; Lamme, 2010; Lamme et al., 2002; Noorman et al., 2025a, 2025b).

For the following analyses, we conducted an omnibus repeated-measures ANOVA with the factors stimulus feature (rotation, collinearity, illusion), masking (unmasked, masked), expectation (expected stimulus in a block, unexpected stimulus in a block), task-relevance (decoded feature is task-relevant, decoded feature is task-irrelevant), and drug (memantine, placebo) on mean classification performance (AUC) per feature. Note that for the factor task-relevance (attention) we decoded the task-relevant feature of the task in a given block (e.g., illusion when focusing on the illusion) and compared it to decoding of that same feature when it was task-irrelevant and participants performed a task on one of the other stimulus features (e.g., illusion when focusing on rotation). The complete results of this omnibus ANOVA are presented in Table 1 and are plotted in **Fig. 3**, together with the EEG decoding time courses, split across all conditions. Full temporal generalization analyses are shown in the Supplementary **Fig. S2**. These matrices reveal consistent generalization patterns across time, indicating that differences in decoding were not driven by shifts in response latency or polarity.

**Figure 3.**
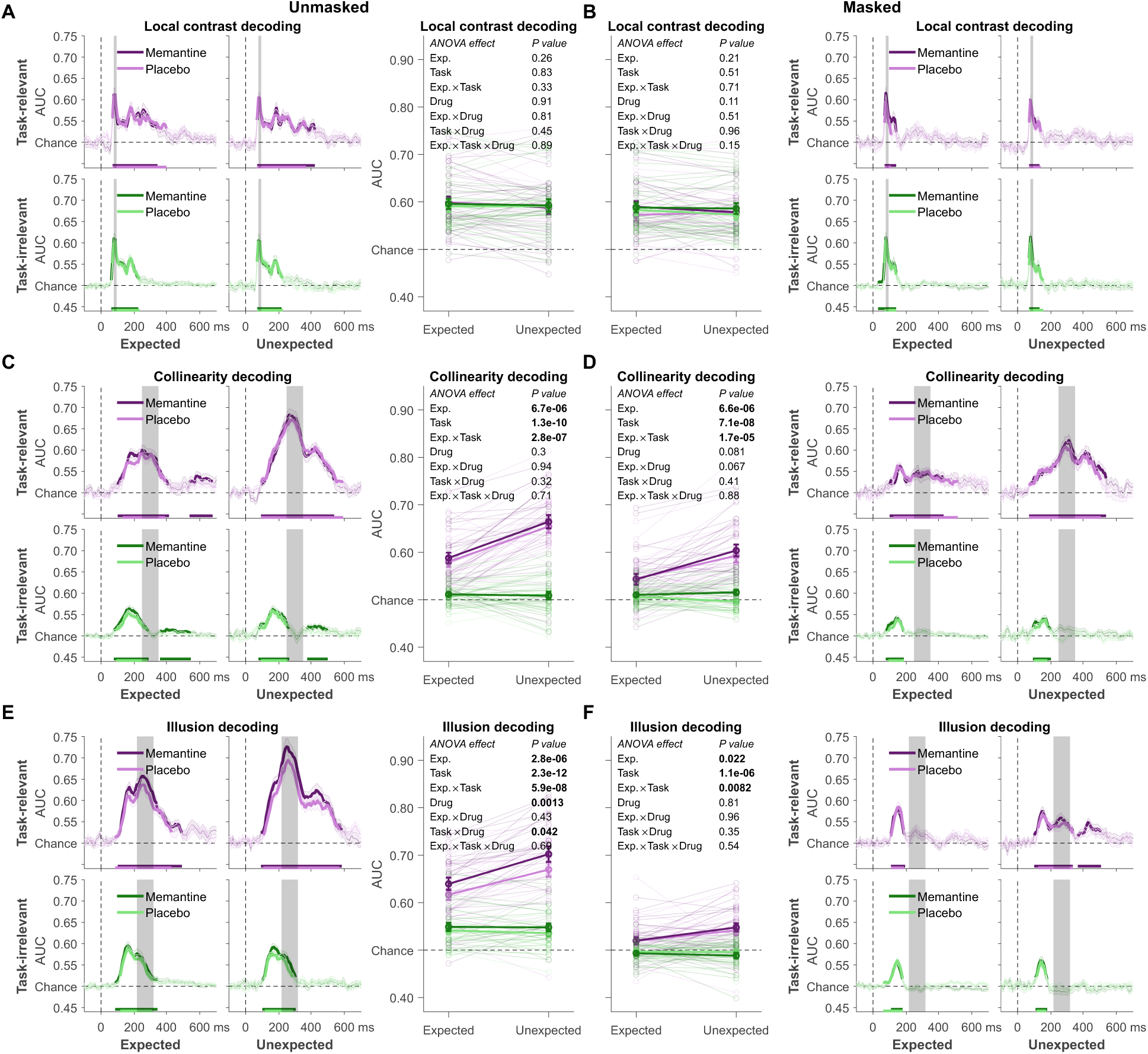
(A-B) Local contrast, (C-D) non-illusory triangle, (E-F) and illusory triangle decoding for all factors in our design. On the left of each panel: mean decoding performance, area under the receiver operating characteristic curve (AUC), over time ± standard error of the mean (SEM). Thick lines represent time-points that differ from chance: P<0.05, cluster-based permutation test. On the right of each panel: For every condition, mean AUC of the decoded feature’s time window. Error bars are mean ± SEM. Individual data points are plotted using low contrast. (A, C, E) Unmasked. (B, D, F) Masked.

**Table 1.** Results of the omnibus repeated-measures ANOVA.

| Effect | F | P | $\eta_p^2$ |
| --- | --- | --- | --- |
| Decoded feature | 8.635 | < .001 | 0.249 |
| Masking | 113.604 | < .001 | 0.814 |
| Expectation | 32.475 | < .001 | 0.555 |
| Task-relevance | 145.433 | < .001 | 0.848 |
| Drug | 5.966 | .022 | 0.187 |
| Decoded feature * Masking | 46.691 | < .001 | 0.642 |
| Decoded feature * Expectation | 28.873 | < .001 | 0.526 |
| Masking * Expectation | 8.666 | .007 | 0.250 |
| Decoded feature * Task-relevance | 59.733 | < .001 | 0.697 |
| Masking * Task-relevance | 62.314 | < .001 | 0.706 |
| Expectation * Task-relevance | 50.963 | < .001 | 0.662 |
| Decoded feature * Drug | 0.335 | .717 | 0.013 |
| Masking * Drug | 0.481 | .494 | 0.018 |
| Expectation * Drug | 0.622 | .438 | 0.023 |
| Task-relevance * Drug | 0.951 | .338 | 0.035 |
| Decoded feature * Masking * Expectation | 1.905 | .159 | 0.068 |
| Decoded feature * Masking * Task-relevance | 25.595 | < .001 | 0.496 |
| Decoded feature * Expectation * Task-relevance | 28.759 | < .001 | 0.525 |
| Masking * Expectation * Task-relevance | 3.772 | .063 | 0.127 |
| Decoded feature * Masking * Drug | 5.682 | .006 | 0.179 |
| Decoded feature * Expectation * Drug | 0.853 | .432 | 0.032 |
| Masking * Expectation * Drug | 0.068 | .797 | 0.003 |
| Decoded feature * Task-relevance * Drug | 1.843 | .169 | 0.066 |
| Masking * Task-relevance * Drug | 1.182 | .287 | 0.043 |
| Expectation * Task-relevance * Drug | 0.005 | .942 | 2.107×10<sup>-4</sup> |
| Decoded feature * Masking * Expectation * Task-relevance | 5.011 | .010 | 0.162 |
| Decoded feature * Masking * Expectation * Drug | 1.024 | .366 | 0.038 |
| Decoded feature * Masking * Task-relevance * Drug | 1.203 | .308 | 0.044 |
| Decoded feature * Expectation * Task-relevance * Drug | 0.790 | .459 | 0.030 |
| Masking * Expectation * Task-relevance * Drug | 0.692 | .413 | 0.026 |
| Decoded feature * Masking * Expectation * Task-relevance * Drug | 0.496 | .612 | 0.019 |

To reduce the complexity of unpacking this five-factorial ANOVA, in the following we report the key results following our main hypotheses. We first focused on questions 1 and 2, i.e., the effects of expectations on the processing of features of different complexity and the hypothesized effects of memantine on expectation and feedback processing, before turning to questions 3 and 4, i.e., the direction and timing of the effects of expectation and attention and the interactive effects of expectation and attention on processing of features of different complexity.

### Expectations selectively affect EEG markers of recurrent processing but leave feedforward processing unaffected

The omnibus ANOVA yielded a main effect of expectation (for statistics from the omnibus ANOVA, see Table 1): decoding was better for unexpected than for expected features. Importantly, the interaction between expectation and decoded feature was significant, and separate ANOVAs (shown in **Fig. 4**) for each feature revealed that expectation only affected EEG markers of lateral (collinearity) and feedback (illusion) processing, but had no significant effect on feedforward processing (local contrast). There was also a significant three-way interaction with masking, indicating that the expectation effects for lateral and feedback processing were weaker in the masked than in the unmasked condition but persisted under strong backward masking (see **Fig. 3**). Finally, a repetition-control analysis (**Fig. S4**) showed that, with the exception of local contrast, repetition of a task-relevant feature reduced decoding (consistent with adaptation, Feuerriegel, 2024; Feuerriegel et al., 2021). For example, when the task was to detect the illusion, illusion decoding was significantly better in non-repeat than in repeat trials. However, for illusion decoding this effect of repetition did not interact with expectation, i.e. both expectation and repetition had additive effects on illusion decoding: expectation effects were of similar magnitude for repeated and non-repeated trials, arguing against a single local adaptation mechanism accounting for expectation effects (Grotheer & Kovács, 2015). When task-irrelevant, neither expectation nor repetition affected decodability.

**Figure 4.**
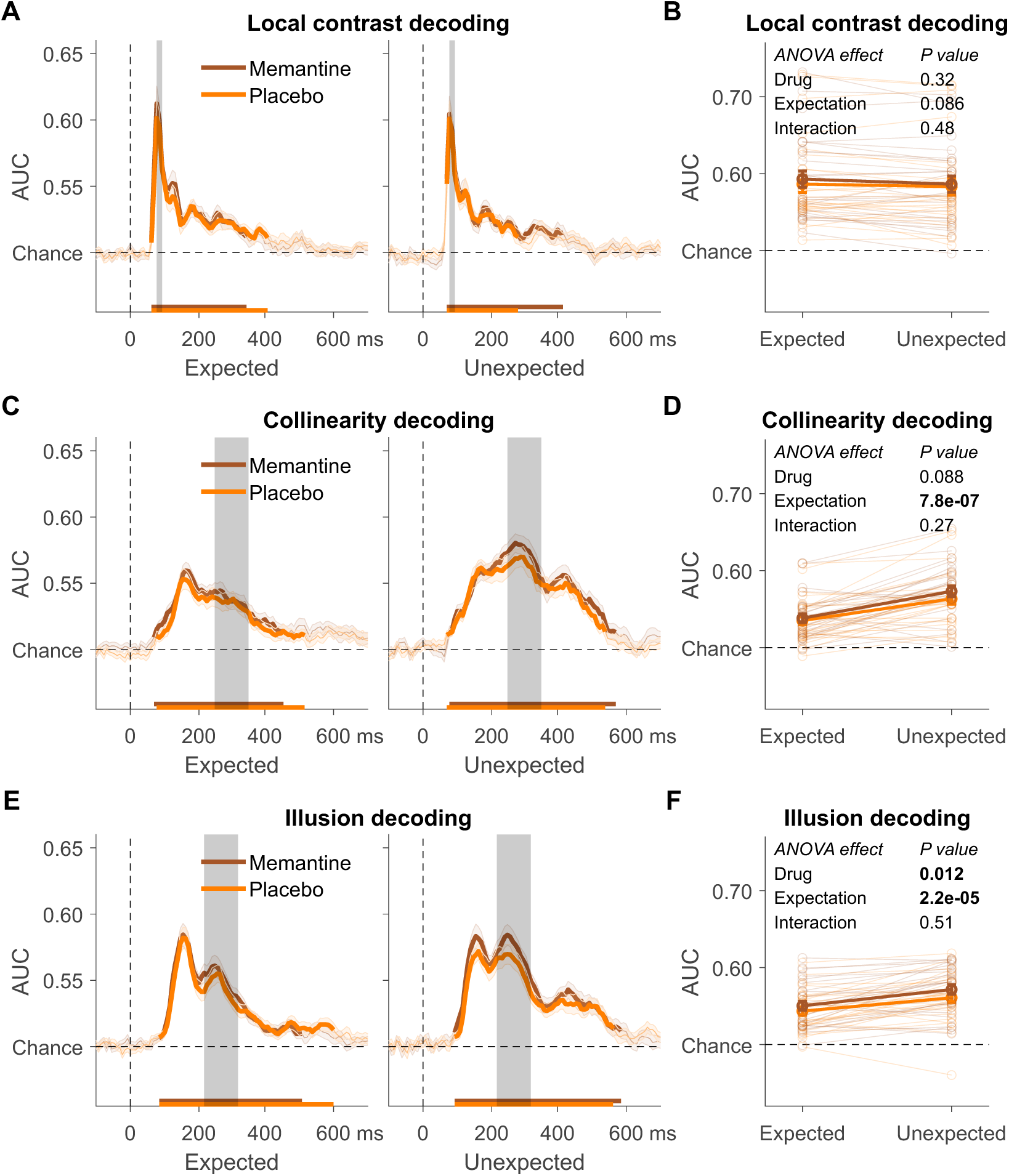
(A-B) Local contrast, (C-D) non-illusory triangle, (E-F) and illusory triangle decoding as a factor of drug and expectation. (A, C, E) Mean decoding performance, area under the receiver operating characteristic curve (AUC), over time ± standard error of the mean (SEM). Thick lines represent time-points that differ from chance: P<0.05, cluster-based permutation test. (B, D, F) For every condition, mean AUC of the decoded feature’s time window. Error bars are mean ± SEM. Individual data points are plotted using low contrast.

### Memantine selectively affects EEG markers of feedback processing but leaves expectations unaffected

Next, we asked whether memantine, in line with its putative effect on NMDA-mediated feedback connections, affected short-term expectations (induced by manipulating stimulus probability) and long-term priors (reflected in illusion processing). The omnibus ANOVA revealed only a main effect of drug and an interaction between masking, feature, and drug (see **Fig. 3** and **Fig. 4**, where we collapsed across masking condition for simplicity). Additional ANOVAs for each of the three different features (shown in **Fig. 4**) revealed, only for the illusion, a main effect of drug. Specifically, memantine *increased* illusion decoding, and this effect was largely specific to unmasked trials (**Fig. 3F**), as we have shown previously (Noorman et al., 2025b). This suggests that memantine affected decoding only when feedback processing was required. Interestingly, memantine did not modulate the neural effects of expectation, also thought to depend on NMDA-mediated feedback processes, as revealed by the absence of any significant interactions between drug and expectation.

### Effects of expectation and attention on EEG markers of lateral and feedback processing

Finally, we focused on the relationship between expectation and attention (task-relevance) and whether they would differ for features of different complexity. The omnibus ANOVA revealed significant two-way interactions with feature for both expectation and attention, as well as significant three-way interactions with masking, which simply reflected somewhat smaller effects in the masked than in the unmasked condition (**Fig. 3**). For simplicity, in **Fig. 5** we again averaged across the unmasked and masked condition. **Fig. 5** shows that the effects of both expectation and attention were specific to collinearity and illusion decoding, while decoding of local contrast was not affected by expectation or attention, indicating that early feedforward processes are relatively immune to these top-down effects. Interestingly, decoding was better for unexpected than for expected stimuli, while attention improved decoding for task-relevant compared to task-irrelevant stimuli.

**Figure 5.**
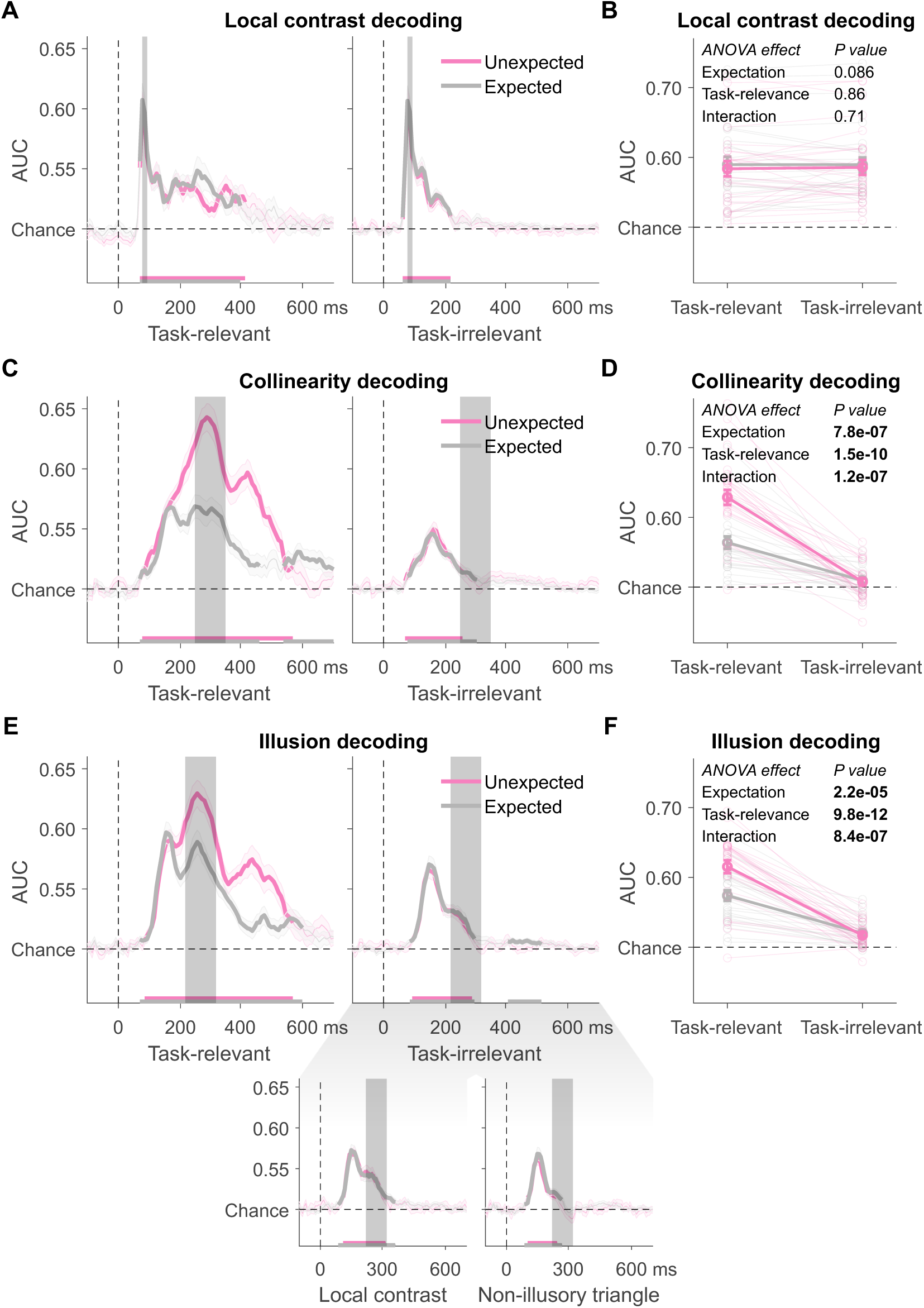
(A-B) Local contrast, (C-D) non-illusory triangle, (E-F) and illusory triangle decoding as a factor of expectation and task-relevance. (A, C, E) Mean decoding performance, area under the receiver operating characteristic curve (AUC), over time ± standard error of the mean (SEM). Thick lines represent time-points that differ from chance: P<0.05, cluster-based permutation test. The inset in (E) shows illusory triangle decoding when the illusion was task-irrelevant and local contrast or the non-illusory triangle were task-relevant, respectively. (B, D, F) For every condition, mean AUC of the decoded feature’s time window. Error bars are mean ± SEM. Individual data points are plotted using low contrast.

For collinearity and illusion decoding, there were significant interactions between expectation and attention, indicating that expectation effects were specific to attended (task-relevant) features, while no reliable expectation effects were observed for task-irrelevant features (**Fig. 5**). For task-irrelevant features, an expectation effect would have been reflected by differences in decoding when those features occurred in blocks where the task-relevant feature was high-probability versus low-probability; such differences were not observed. Because task-irrelevant features were always equiprobable and decoded using within-block presence-absence contrasts, the absence of an expectation effect indicates that expectation-related modulation did not generalize beyond the feature dimension for which expectations were explicitly induced.

Turning to the timing of the effects of expectation and attention, analyses of decoding time courses, averaged across all features, revealed a significant effect of attention beginning 133 ms after stimulus onset, and a significant effect of expectation beginning 188 ms after stimulus onset (**Fig. 6A**). Thus, the rise of the attention effect was significant 55 ms earlier than the rise of the expectation effect, and 141 ms after stimulus onset, the attention effect was significantly larger than the expectation effect (**Fig. 6B**). Note that because cluster-based permutation tests do not provide precise latency estimates (Sassenhagen & Draschkow, 2019), our conclusions primarily concern the relative ordering of effects, i.e. attention preceding expectation, rather than exact absolute onset times.

**Figure 6.**
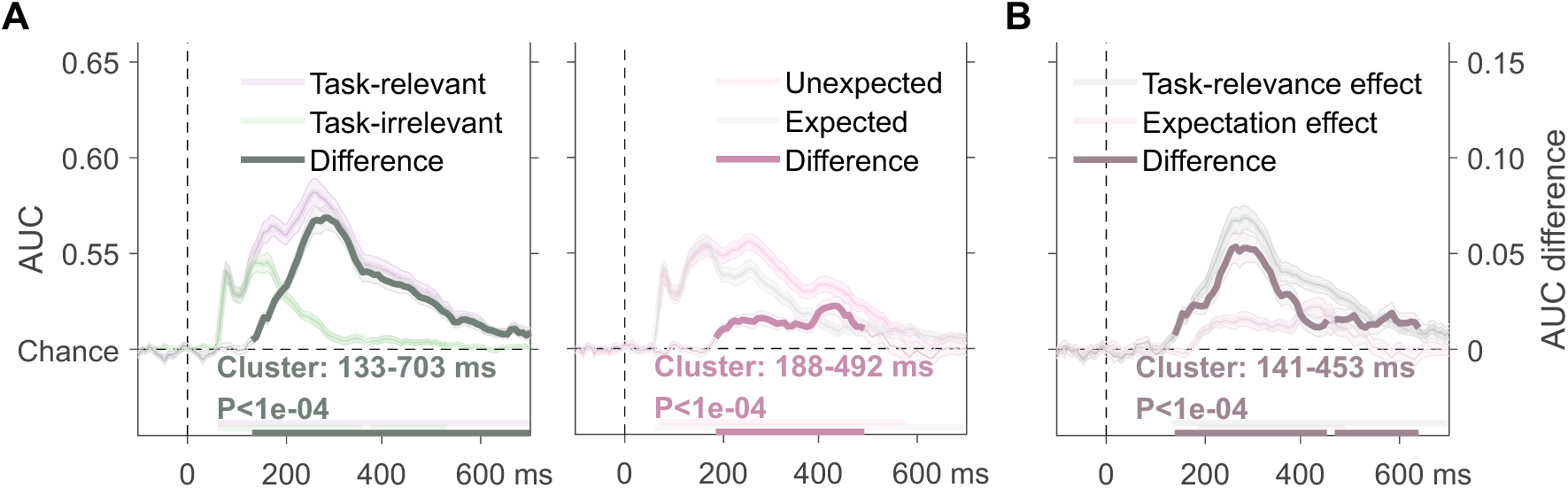
Decoding averaged across all features as a function of (A) the attention (task-relevance) and expectation effects and (B) the difference between them. Mean decoding performance, area under the receiver operating characteristic curve (AUC), over time ± standard error of the mean (SEM). Thick (horizontal) lines represent time-points that differ from chance for the experimental conditions, or from 0 for the differences: P<0.05, cluster-based permutation test.

Further, the difference in decoding between task-relevant and task-irrelevant features was not an artifact of the training procedure. Classifiers trained on localizer data containing only task-relevant trials, only task-irrelevant trials, or both yielded highly comparable results, with consistently stronger decoding for task-relevant than for task-irrelevant features (**Fig. S3**). This confirms that the effect of attention reflects genuine differences in feature processing rather than a bias introduced by classifier training.

## Discussion

This study tested the effects of expectation on feedforward vs. recurrent neural processing of stimulus features of different complexity, their relation to attention, and their putative NMDA-receptor mediated biochemical underpinnings. We found that expectations about stimulus probability had distinct effects at early vs. later stages of processing: while EEG decoding of complex features involving recurrent processing (collinearity, illusion) was enhanced when these features were unexpected, decoding of local contrast, relying on feedforward processing, was not affected by expectation. Similarly, attention (task-relevance) enhanced decoding of collinearity and of the illusion, but had no effect on early decoding of local contrast. Importantly, expectation effects were observed only for task-relevant features, whereas no reliable expectation effects were found for task-irrelevant features. Effects of expectation persisted under masking, were absent for local-contrast decoding, and were mostly additive with feature repetition, supporting the conclusion that our base-rate manipulation induced genuine expectation-related modulation rather than low-level adaptation. Finally, while the NMDA antagonist memantine did not affect the neural effects of expectation, this manipulation did affect decoding of the Kanizsa illusion. This indicates that feedback-modulating effects were specific to long-term priors about the spatial arrangement of stimuli (and the associated perception of a Kanizsa triangle) but did not extend to expectations based on shorter-term manipulations of stimulus probability.

The first question we asked was whether expectation affects all processing stages similarly, as suggested by some theoretical models (Friston, 2005; Lee & Mumford, 2003). Instead, and in line with recent EEG findings (Alilović et al., 2019; Rungratsameetaweemana et al., 2018; Rungratsameetaweemana & Serences, 2019), we found that expectations primarily modulated decoding of more complex features associated with recurrent processing (collinearity and especially illusion). This selectivity to later processing stages was also found for other manipulations of recurrent processing: like expectation, attention (task-relevance) did not influence the most low-level feature (local contrast); and while masking did have a significant, albeit weak, effect on local contrast decoding, its impact on decoding of collinearity and illusion was relatively larger. Overall, this pattern of results is in line with recurrence being progressively more important for later processing stages engaged in the perception of more complex features (Gilbert & Li, 2013; Lamme & Roelfsema, 2000). However, it stands in apparent contradiction to fMRI findings of expectation effects already in primary sensory areas (Alink et al., 2010; Kok, Jehee, et al., 2012; Murray et al., 2002; Richter & de Lange, 2019; Richter et al., 2017). One reason for this discrepancy might be the temporal resolution of fMRI, making it difficult to distinguish early sensory processing from later downstream effects on early sensory areas (Nienborg et al., 2012; Wilming et al., 2020). Although our design did not permit reliable single-trial correlations between decoding strength and behavior, the parallel pattern of behavioral and neural effects across conditions supports the interpretation that our decoding reflects perceptual effects, mediated by recurrent processing rather than low-level feedforward processing.

Our findings also underscore the importance of NMDA receptor-mediated feedback mechanisms in perceptual integration. The administration of memantine, an NMDA receptor antagonist, selectively enhanced decoding of the Kanizsa illusion but did not affect decoding of local contrast or collinearity. Although a reduction in decoding might have been expected given memantine’s NMDA-antagonistic profile, the observed enhancement is consistent with our earlier report; we refer interested readers to the in-depth treatment in Noorman et al., 2025b. Importantly however, memantine did not modulate the neural effects of short-term expectations, suggesting a dissociation between the mechanisms underlying the implementation of long-term priors (such as those involved in processing illusory figures) and those driving the modulation of sensory processing by short-term expectations induced by task demands. It is possible that different neurochemical systems, for example GABAergic or dopaminergic pathways (Bastos et al., 2012) or different aspects of the NMDA receptor’s role are involved in these distinct processes (Meuwese et al., 2013; Thiele & Bellgrove, 2018).

While expectation and attention showed similar selectivity to later processing stages, the direction of their effects on EEG decoding was opposite: decoding of expected features was reduced, while decoding of attended (task-relevant) features was enhanced. One possible explanation for this dissociation could have been that unexpected stimuli captured attention (Alink & Blank, 2021). However, this account is not supported by the present results. If unexpected stimuli had globally attracted attention, expectation effects should also have been observed for task-irrelevant features. Instead, expectation-related modulation was confined to task-relevant features and absent for task-irrelevant features. This pattern indicates that expectation effects in the present paradigm did not arise from stimulus-driven attentional capture, but were selectively expressed, in a top-down manner, for features that were behaviorally relevant.

In line with opposing process theory (Press et al., 2020), such expectation-related suppression emerged at later processing stages associated with recurrent processing (collinearity and illusion), and with a delayed onset relative to attentional enhancement. In this view, expectations may “dampen” neural responses to predictable inputs, selectively reducing information about expected (and therefore less informative) stimulus features (Press et al., 2020; Richter et al., 2017, 2022). This expectation-driven cancellation mechanism is thought to come into play when an unexpected input is strong (i.e., has low sensory noise, like the high-contrast Kanizsa stimulus used here), rendering it highly surprising, upweighting processing of unexpected inputs relatively “late” in time (∼150–200 ms after stimulus presentation), following an initial bias towards expected inputs. Our findings are consistent with this theory: expectation-related suppression occurred only for later processing stages representing collinearity and the illusion, and was evident only 188 ms after stimulus onset, later than attention-related enhancement, which began already 133 ms after stimulus onset. Furthermore, while the behavioral results revealed a response bias to indicate feature presence for expected inputs, perceptual sensitivity was greater for unexpected inputs, as predicted by opposing process theory. Importantly, because the three stimulus features were fully orthogonalized and balanced across all conditions, each varied independently of the others, ensuring that these expectation effects on task-irrelevant features cannot be attributed to higher-order feature interactions.

The finding that expectation effects were limited to task-relevant features places important constraints on claims about the relationship between expectation and attention. On the one hand, our results are compatible with proposals that expectation and attention exert dissociable influences on neural processing (Summerfield & de Lange, 2014; Summerfield & Egner, 2009). On the other hand, they are also consistent with studies suggesting that attention can gate, constrain, or even reverse the expression of expectation effects (Kok, Rahnev, et al., 2012; Larsson & Smith, 2012; Richter & de Lange, 2019). In those studies (e.g., Richter & de Lange, 2019; Richter et al., 2017), expectation effects were abolished when attention was fully diverted away from the stimulus, whereas they remained intact when stimulus regularities were task-irrelevant but attention was not explicitly withdrawn. In our present design, expectations were manipulated only for task-relevant features, and expectation effects were observed only along that attended dimension. This pattern suggests that, at least for short-term expectations based on stimulus probability, attentional engagement with a feature dimension may be a prerequisite for expectation-related modulation to emerge.

Taken together, the present findings suggest that top-down influences on perception are not uniform across processing stages or cognitive manipulations. Earlier, predominantly feedforward stages of visual processing were comparatively less affected by both expectation and attention, whereas later stages that rely more strongly on recurrent (lateral and feedback) processing were modulated by both, albeit in different ways. Expectation-related suppression was selective to task-relevant features and did not generalize to task-irrelevant features, while attentional enhancement robustly increased decoding for task-relevant features. The absence of an effect of NMDA receptor blockade on expectation-related modulation further indicates that short-term, probability-based expectations rely on mechanisms that are at least partially distinct from those supporting long-term perceptual priors, such as those underlying illusory contour perception. By combining time-resolved EEG decoding with manipulations of expectation, task relevance, feature complexity, and NMDA receptor function, the present approach provides a principled framework for dissociating distinct forms of top-down influence on sensory processing. This integration of causal pharmacological manipulation with temporally resolved neural markers offers a powerful methodological avenue for testing mechanistic accounts of perceptual inference and recurrent processing in the human brain.

