## Supplemental Material for "Effects of expectation, attention, and NMDA receptor blockade on feedforward and feedback processing"

This research was supported by a grant from the H2020 European Research Council (ERC STG 715605, SVG).

### Supplementary Material

**Table of Contents**

Figure S1. Behavioral results for drug and placebo conditions.

Figure S2. Temporal generalization matrices.

Figure S3. Decoding of task relevance as a function of training data set.

Figure S4. Repetition-control analysis.

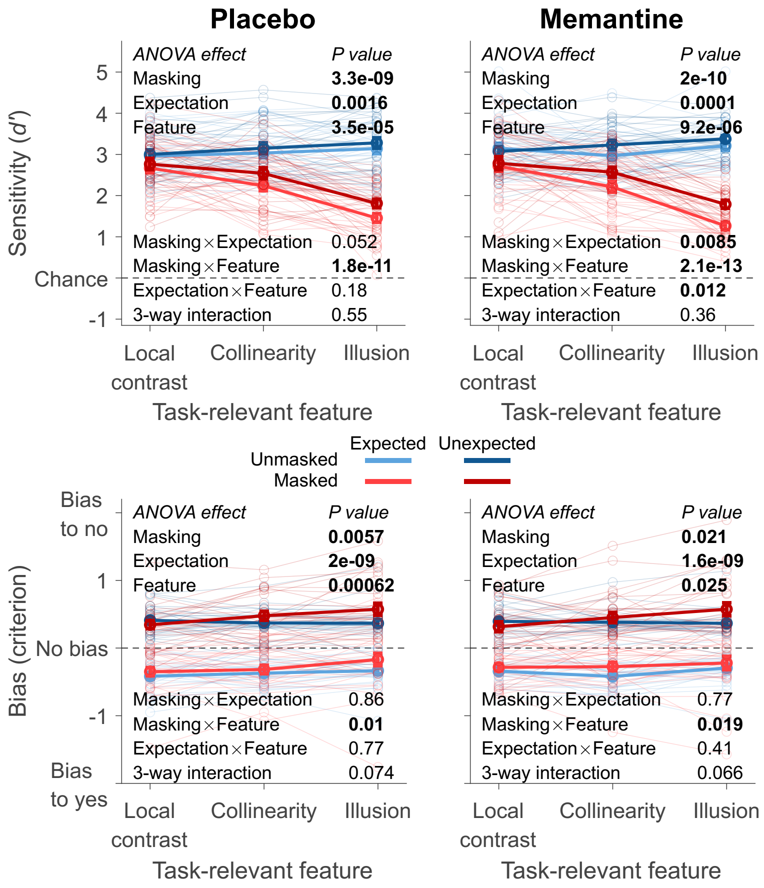


***Figure S1. Behavioral results for drug and placebo conditions.*** *Perceptual performance and bias related to participants' ability to detect the task-relevant (attended) stimulus dimension. Error bars are mean ± standard error of the mean. Individual data points are plotted using low contrast.*

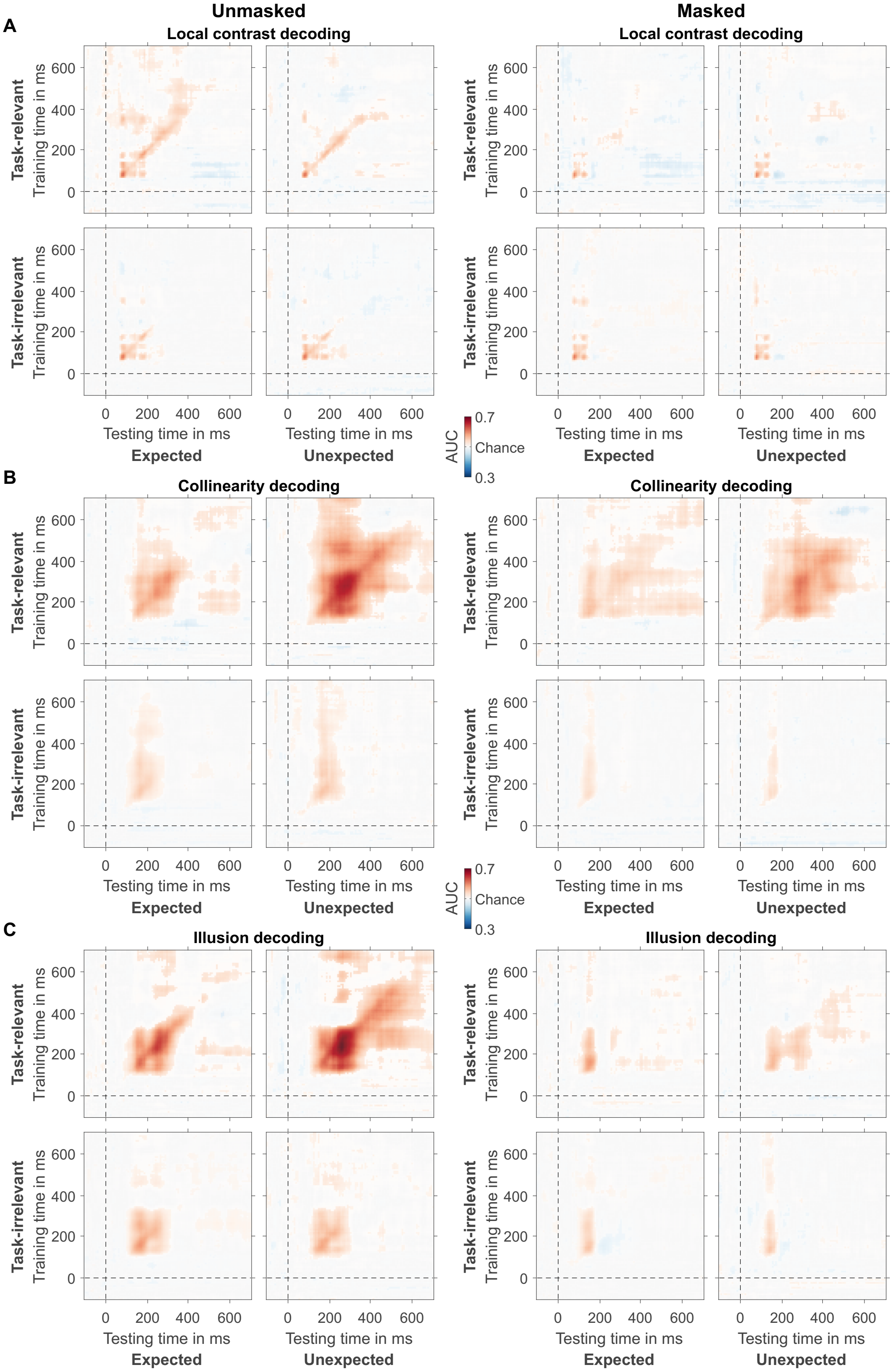


***Figure S2. Temporal generalization matrices.*** *Panels show mean decoding performance (AUC) for different combinations of training and testing times, separately for (A) local contrast,*

*(B) collinearity, and (C) illusion decoding. Left panels: unmasked, right panels: masked. Sub-panels show all factorial combinations of expectation and task-relevance.*

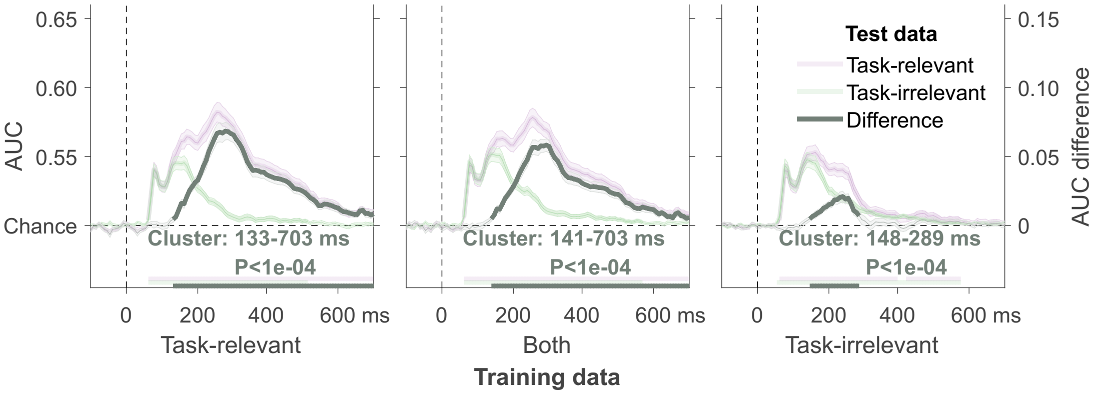


***Figure S3. Decoding as a function of training data set.*** *Classifiers were trained on localizer data containing only task-relevant features, both task-relevant and task-irrelevant features and task-irrelevant features only (shown in the separate panels from left to right). Decoding averaged across all features for task-relevant and task-irrelevant test stimuli and the difference between them. Mean decoding performance, area under the receiver operating characteristic curve (AUC), over time ± standard error of the mean (SEM). Thick horizontal lines represent time-points that differ from chance for the experimental conditions (purple and green lines), or from 0 for the differences in grey: P<0.05, cluster-based permutation test.*

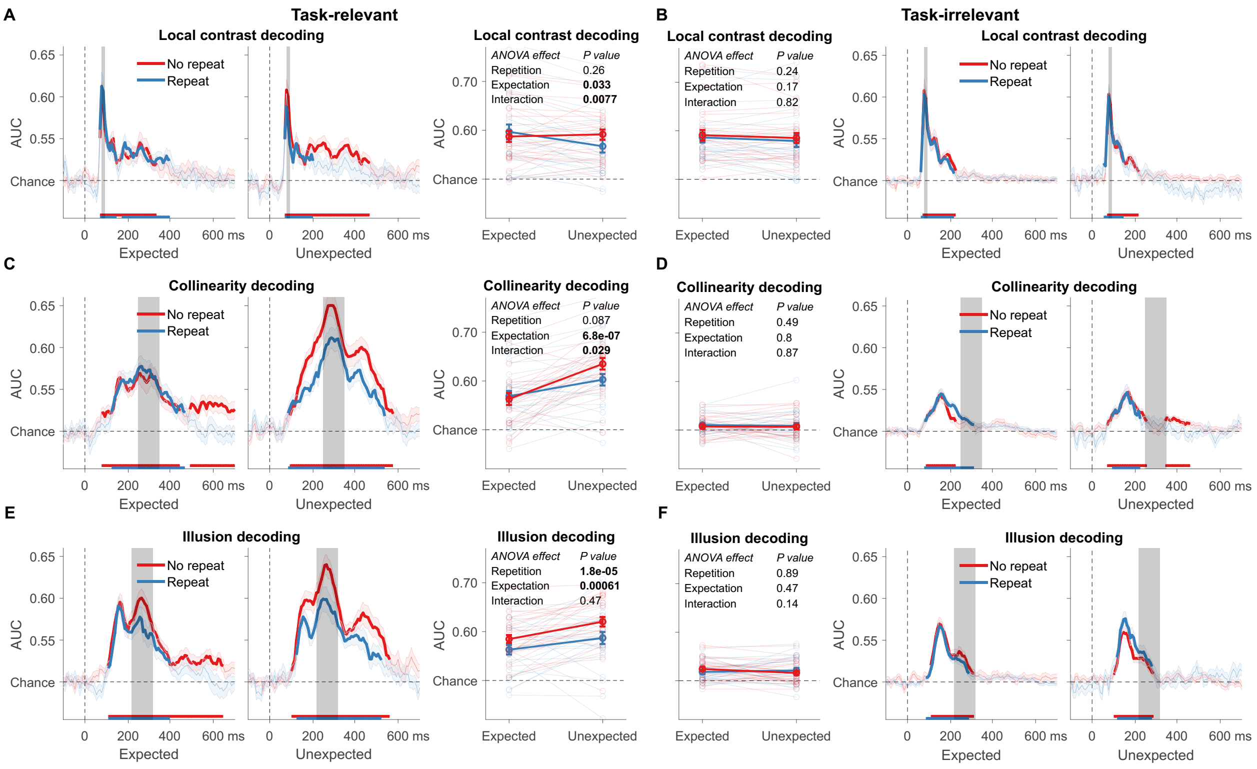


***Figure S4. Repetition-control analysis.*** *Control analyses tested whether expectation effects reflect adaptation related to feature repetition. We split trials into repeated versus non-repeated (same feature as previous trial vs. not). (A-B, G-H) Local contrast, (C-D, I-J) non-illusory triangle, (E-F, K-L) and illusory triangle decoding separately for expected and unexpected conditions. The left panels (A–F) show decoding for task-relevant features; the right panels (G-L) show decoding for task-irrelevant features. For this analysis we collapsed across the factors masking and drug. On the left of each panel: mean decoding performance, area under the receiver operating characteristic curve (AUC), over time ± standard error of the mean (SEM). Thick lines represent time-points that differ from chance: P<0.05, cluster-based permutation test. On the right of each panel: For every condition, mean AUC of the decoded feature's time window. Error bars are mean ± SEM. Individual data points are plotted using low contrast.*
